# Advancing Microbial Comparative Genomics Through Tumor-Normal Inspired Framework

**DOI:** 10.1101/2025.02.05.636605

**Authors:** Junjie Yang, Sheng Yang

**Author notes:** **Address:** 300 Fenglin Road, Shanghai, 200032, China.

## Abstract

Comparative genomics has emerged as a pivotal methodology for elucidating genetic variations in microbial studies. However, conventional analytical approaches for diploid and polyploid microorganisms have demonstrated limited efficacy in discriminating new mutations from background heterozygosity.

This study presents an innovative microbial comparative genomics framework adapted from tumor-normal sequencing methodology. Our approach establishes the original strain as the “normal sample” and the derived strain as the “tumor sample”, enabling precise identification of new mutations (“somatic variants”) while filtering pre-existing heterozygous sites (“germline variations”). The analytical pipeline also includes assessment of loss-of-heterozygosity (LOH) events and genome-wide detection of copy number variations (CNVs) with resolution to identify both regional CNV and whole-chromosome aneuploidy through integrated CNV and variant allele frequency analysis. We validated this framework using diploid *Saccharomyces cerevisiae* strains before successfully extending its application to *Kluyveromyces marxianus*, *Candida* spp., and *Hortaea werneckii*, encompassing haploid, diploid, polyploid, and aneuploid states. The methodology revealed previously undetected variations across experimental evolution studies, demonstrating superior resolution compared to conventional approaches. This adaptable platform establishes a new paradigm for microbial genome studies, particularly for organisms with diploid or polyploid states where traditional comparative genomics methods prove inadequate.

## Introduction

Comparative genomics provides a powerful lens to investigate genetic variations arising during microbial adaptation, including laboratory-driven evolution, mutagenesis, and spontaneous mutation processes. A pivotal challenge lies in distinguishing relevant mutations underlying phenotypic adaptation from background genetic noise--natural heterozygous loci.

Conventional analytical approaches for diploid microorganisms, including reference-based variant detection and de novo assembly pipelines, exhibit fundamental limitations in addressing this challenge. While differential variant calling between progenitor and evolved strains theoretically enables identification of new mutations, substantial natural heterozygosity in diploid genomes (typically ∼10,000 heterozygous loci in *Saccharomyces cerevisiae* native strains) introduces systematic artifacts. Notably, even modest false-positive detection rates (e.g., 1%) generate ∼100 spurious variants, effectively obscuring genuine adaptive mutations. Assembly-based methodologies similarly fail to resolve this issue due to inherent difficulties in resolving heterozygous regions. These analytical constraints impose significant burdens on downstream functional validation and create persistent bottlenecks in establishing robust genotype-phenotype correlations.

To address these limitations, we present an analytical framework that adapts tumor-normal sequencing paradigms(1)—originally developed for somatic mutation detection in cancer genomics—to microbial studies. In our prior work applying this approach to evolved diploid *S. cerevisiae* strains with enhanced lignocellulosic hydrolysate conversion capabilities, we focused primarily on metabolic and physiological characterization while providing limited methodological details. (2) This study provides a comprehensive exposition of the analytical pipeline. To demonstrate the versatility, we extend its application beyond *S. cerevisiae* to diverse microbial systems spanning multiple ploidy states, including *Kluyveromyces marxianus*, *Candida* spp. and *Hortaea werneckii.* This systematic implementation establishes a generalized framework for mutation discovery.

## Results

### Tumor-Normal Sequencing Framework for Microbial Comparative Genomics

This analytical framework adapts tumor-normal sequencing principles from cancer genomics to microbial experimental evolution. The progenitor strain is designated as the “normal sample”, while the evolved strain serves as the “tumor sample”, enabling systematic identification of *somatic variations*(novel genetic changes acquired during adaptation) while filtering *germline variations* (pre-existing heterozygous loci, reference genome errors, or assembly artifacts). This paired approach minimizes false positives arising from natural genetic diversity and technical noise. This analytical framework is schematically summarized in Figure 1.

**Figure 1.**
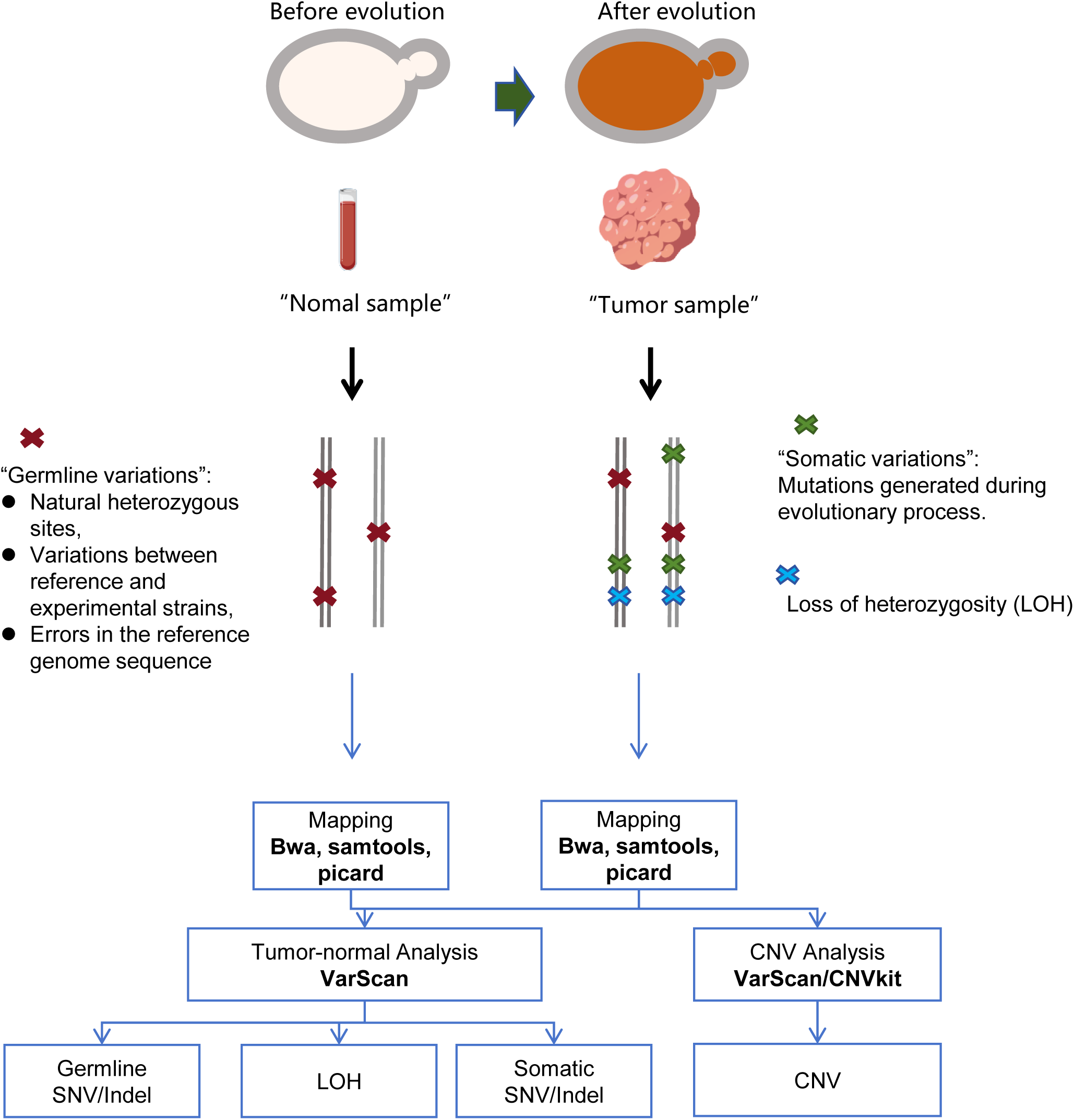
Schematic workflow for microbial comparative genome analysis using the tumor-normal sequencing conceptual framework.

Integrative visualization employs a chromosome-coordinate scatter plot to resolve variant classification: somatic mutations (novel alleles in evolved strains), germline polymorphisms (pre-existing heterozygous loci, etc.), and LOH events (complete allele fixation) are demarcated by their variant allele frequencies (VAF) and genomic distribution (Figure 2).

**Figure 2.**
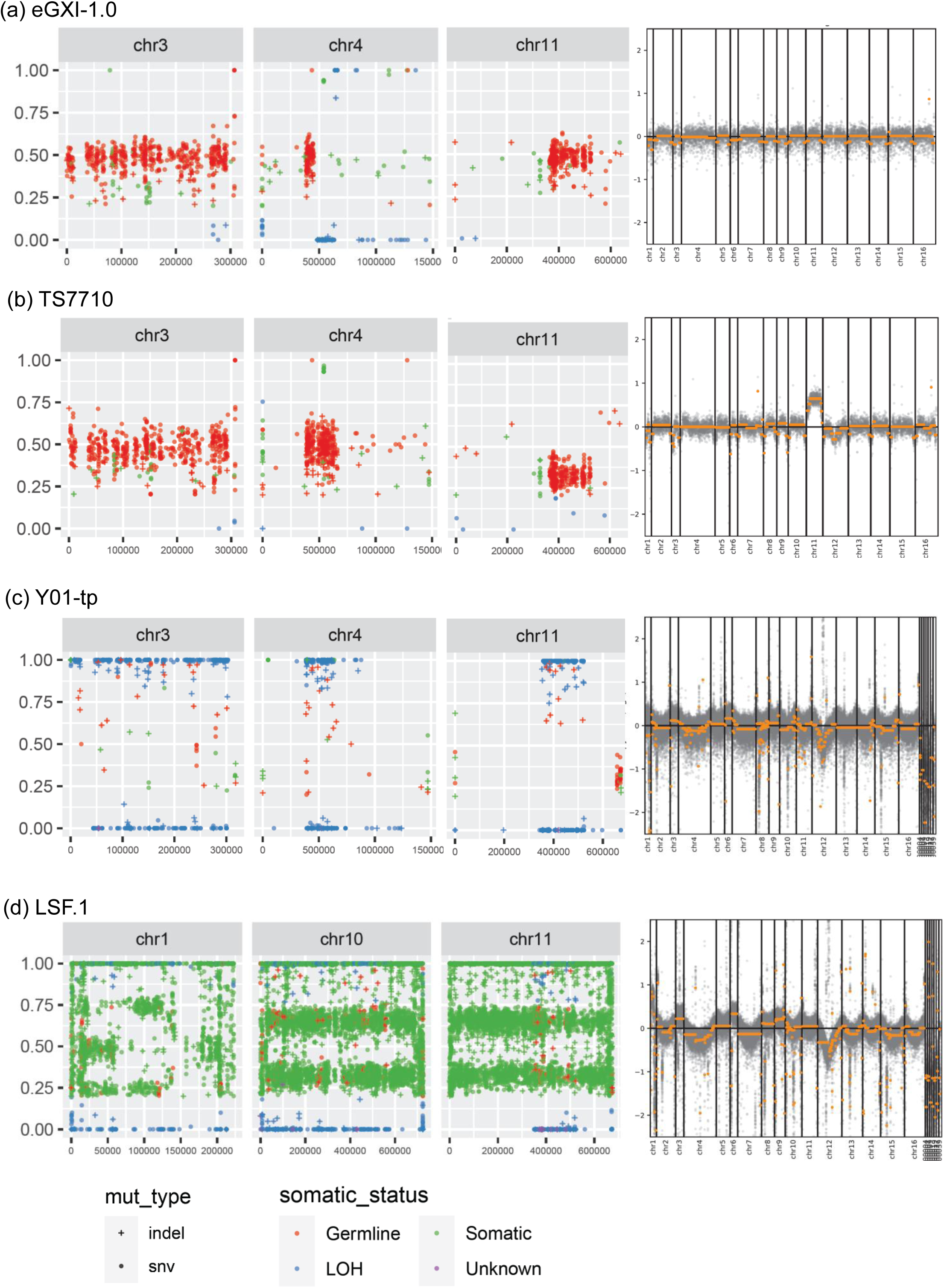
Genome-wide distribution profiles of genetic variations (left) and copy number variations (CNVs, right). (a) *Saccharomyces cerevisiae* eGXI-1.0: diploid strain exhibiting somatic variations, germline variations, and loss of heterozygosity (LOH) on chromosome 10. (b) CIBT7710: diploid strain displaying somatic variations, germline variations, LOH on chromosome 10, and CNV on chromosome 11 leading to aneuploidy 2n+1(XI). (c) Haploid strain Y01-tp: majority of variations detected as homozygous, including homozygous somatic variations and LOH. (d) LSF.1: numerous somatic variations identified when using Ethanol Red as normal sample, reflecting substantial genetic divergence between them. The copies of most chromosomes should be 3 and the copies of chromosome 1 should be 4, leading to aneuploidy 3n+1(I).

Copy number variation (CNV) analysis detects both segmental duplications/deletions and whole-chromosome aneuploidy. (Figure 2)

### Validation and Case Studies

To demonstrate universal applicability, we present several Case Studies: Case Study 1 is using our recent publication samples about *S. cerevisiae* adaptive laboratory evolution as mentioned above. (2) Other Case Studies are using samples reported in recent articles, with new discoveries made by our analysis. (Table S2)

### Case Study 1

Specifically, we evolved a diploid *S. cerevisiae* strain that overexpresses xylose catabolism genes under conditions of mixed xylose/glucose medium supplemented with sodium salts. The fully evolved yeast strains were obtained after 182 transfer demonstrated efficient conversion of glucose and xylose in lignocellulose hydrolysates to ethanol, highlighting their potential for industrial applications. To uncover the mechanical behind the complex phenotypes of high resistance to inhibitors and efficient conversion of glucose and xylose in lignocellulose hydrolysates to ethanol, accurately detecting new variations becomes a key step. The tumor-normal sequencing concept for microbial comparative genome analysis were performed. (2)

In the final evolved strain eGXI-1.0 after 182 transfers with improved efficient conversion of glucose and xylose in lignocellulose hydrolysates to ethanol, 288 high confidence somatic variations (231SNV + 57Indel) were detected. (Figure 2a) Meanwhile thousands of germline variations were detected, most of which are native pre-existing heterozygous sites and will not be researched in our experimental research.

Among the somatic variations – new mutations during the evolution process, mutations in *TRK1*, *CCC1* etc. were identified as key mutations and experimentally confirmed in our subsequent study. (2)

We also detected 502 high confidence LOH variations (377SNV + 25Indel) in this evolved strain. With the scatter plot for presenting the overall distribution of variations, large LOH in chromosome 4 and chromosome 10 were shown. As the cassette of *xylA* (xylose isomerase) integrated into one homologous chromosome of chromosome 4, it is speculated that the LOH of chromosome 4 results in the chromosome with *xylA* replaced the homologous chromosome without *xylA*, helping the copy number application of this xylose metabolism gene.

Meanwhile, CNV analysis is performed. Large CNV, including chromosomal aneuploidy was not detected in eGXI-1.0. (Figure 1). The VAF, approximately 50% (heterozygous variations) or 100% (homozygous variations and LOHs), confirmed diploidy of this strain.

We also conducted the same analysis on the strain CIBT7710 which was isolated from the cell mixture after the 60 transfers of the evolution process. High quality somatic variations (148 SNV + 54 Indel) were detected. (Figure S2) LOH in chromosome 4 was not detected in this strain, indicating that the LOH event for increasing the copy of *xylA* happened in later generations. (Figure 2b)

Copy number amplification of whole chromosome 11 was detected in CIBT7710. Meanwhile the VAF, approximately 33 or 67%, confirmed that chromosome 11 should amplify to 3 copies. Therefore, the ploidy of this strain should be aneuploidy 2n+1 (XI). (Figure 2b)

### Case Study 2

A method named the Genomewide Evolution-based CRISPR/Cas with Donor-free (GEbCD) system was used for editing industrial diploid *S. cerevisiae* strains SISc, by which strain G4-72 with strong robustness and higher productivity was obtained. It is reported that 1711 resulting-strain-only variations (1396 SNV + 315 Indel) were detected in G4-72. Also, it is reported that G4-72 displayed a 1.5-fold increase in the relative copy number of chromosome 9 compared with SISc-Δrad52, suggesting the occurrence of an additional chromosome 9 copy. (3)

The chassis strain of this study is another industrial ethanol yeast used in China and with high genetic relationship with the original strain in Case Study 1, so analysis was performed using our original strain AQ as normal sample. The results showed that 328 (267SNV+ 61 Indel) and 1270 (1135 SNV + 145 Indel) somatic variations were detected respectively in the beginning strain SISc and the resulting strain G4-72 of GEbCD operation (Figure S1). The resulting-strain-only high confidence variations are only 1090, which is less than the previous report. We speculate that some of the mutations reported in this article may be natural heterozygous sites but mistakenly identified as new mutations. Therefore, the effectiveness of this GEbCD system might need to be reevaluated and perhaps be more accurate than the previous report. Besides, our analysis results confirmed the applications in chromosome 9. (Figure S1)

### Case Study 3

Using this process, we analyzed genome sequencing data released from several studies on *S. cerevisiae* strain Ethanol Red (from Lesaffre), another widely used industrial ethanol yeast.

An evolved industrial *S. cerevisiae* strain ISO12 with improved tolerance to grow and ferment in the presence of lignocellulose-derived inhibitors at high temperature (39 °C) was reported. Through whole-genome sequencing and variant calling, a high number of strain-unique SNPs and INDELs were identified (1472 total, 306 non-synonymous). (4) LOH analysis was not performed in this report. Three ORFs exhibited a CNV higher than threefold, including *ENA*, *PHO12* and *FLO1*. (4)

Our analysis identified 603 high confidence somatic variations (497SNV+106Indel) much less than the previous report. Only 187 mutations (upstream of the coding region + non-synonymous in ORF) were considered potentially with higher importance for phenotype. Meanwhile, our analysis also identified LOH in chromosome 1, chromosome 10, chromosome 13, chromosome 9, and several large fragment amplifications in chromosome 4 and chromosome 14. So, mutations, LOH and CNV may play a combined role in phenotypic changes in strain ISO12. (Figure S1)

ScY01 is a strain derived from Ethanol Red through adaptive evolution at high temperature.(5) One osmotolerant strain ScY001T and one thermotolerant strain ScY033T were selected from the mutant library generated by TALENs-assisted multiplex editing (TAME) at corresponding stress conditions and exhibited 1.2-fold to 1.3-fold increases of fermentation capacities, respectively.(6) By comparing with the reference genome of strain S228c, the different SNPs and InDels were detected in ScY001T versus ScY01 (1705 + 514) as well as ScY033T versus ScY01 (1718 + 498). (5, 6) LOH analysis was not performed in this report. CNVs of the selected stress-tolerant strains were assessed for both nuclear genomic DNA and mitochondrial genomic DNA (mtDNA). Ampliffcation-type and deletion-type CNVs were equally distributed across nuclear chromosomes while the majority of CNVs were amplification-type on mtDNA. But no detailed analysis about CNVs was shown. (5, 6)

Our analysis identified 203 and 206 high confidence somatic variations (130 CNV+73 Indel, 128SNV + 78 Indel) in ScY001T and ScY033T, much less than the previous report. Meanwhile, amplification of chromosome 9 causing aneuploidy was detected in all these 3 strains, same as G4-72 in Case Study 2. The function of chromosome 9 amplification is worth further research. Large LOH were detected for all the 3 strains in several chromosomes but with different patterns. (Figure S1) Mutations, LOH and CNV may play a combined role in phenotypic changes in these strains.

We also applied this analytical method to a haploid strain Y01-tp, which is an haploid segregant from ScY01.(7) The results showed that almost all of the variations were homozygous variations (LOH and homozygous somatic variation), which is consistent with the characteristics of haploid. (Figure 2c) This result provides a typical example for haploid (or highly homozygous) strains of this analysis.

### Case Study 4

We analyzed another active dry yeast(ADY) strain LSF.1, which is reported to be from Lesaffre, but lacks detailed information.(8) With Ethanol Red as normal and reference genome sequence, dozens of thousands of somatic cell variations(93083 SNV+ 6290Indel) were detected in LSF.1, indicating that this strain was not derived from Ethanol Red or close to Ethanol Red (Figure 2d). This strain should be the ADY in fields other than ethanol, probably in mantou or bread.

This result provides a typical example that the final strain (tumor sample) and the initial strain (normal sample) do not come from the same ancestral. For instance, if unexpected contamination happened in evolution experiments, the resulting strains might show similar results as this example.

Meanwhile, the VAF on most chromosomes are around 33% or 67%, but the VAF on chromosome 1 are around 25%, 50%, and 75%. The CNV results show an application of chromosome 1. (Figure 1)The copies of most chromosomes should be 3 and the copies of chromosome 1 should be 4, so ploidy of this strain should be aneuploidy 3n+1(I), consistent with previous reports by Flow Cytometry (8).

### Case Study 5

We also used this analysis method to analyze other yeasts, first applying it to *Kluyveromyces marxianus*, an emerging yeast for food and biotechnology applications.

By adaptive evolution in 6% (v/v) ethanol for 100-day evolution from wild-type K. marxianus FIM1, the evolved strain KM-100d was obtained with ethanol tolerance increased up to 10% (v/v). SNPs located in the coding region of *SAN1*, *YAP1* were detected and predicted related to ethanol tolerance in the previous report.(9)

Our analysis detected 41 somatic variations (32 SNV+9 Indel) inducing only 4 high confidence variations (3 SNV+1 Indel). SNVs located in the *SAN1* and *YAP1* detected in our results correspond to the previous reports. We detected a ∼100kb amplification region, located at CP015056.1 (chromosome 3) (1476415∼1571769), harboring several transcription related genes, such as *IMP2* (sugar utilization regulatory protein IMP2), *RRD1* (serine/threonine-protein phosphatase 2A activator 1), *SLN1* (osmosensing histidine protein kinase SLN1). (Figure S2) We speculate the dose of these genes may be associated with ethanol tolerance.

### Case Study 6

*Candida tropicalis* is a nonconventional yeast with medical and industrial significance. As a xylose-fermenting yeast, it has the potential for converting cellulosic biomass to ethanol. *C. tropicalis* X-17.2b is an evolved strain with increased tolerance to ethanol, furfural and hydroxymethyl furfural at high temperatures and improvement in glucose/xylose fermentation ability at high sugar concentrations. But in the previous report, mutation points in X-17.2b were not been identified. (10)

Our analysis detected 373 high confidence somatic variations (163 SNV+210 Indel), in theory equivalent to new mutations during evolution. We manually picked up several mutations including the mutations in the upstream region of *NGT1*(High-affinity glucose transporter 1) and *GSF2* (Glucose signaling factor 2), which should be related to the transcriptional level changes of *NGT1* and *GSF2* (10). More genes regulated by GSF2 might affect the phenotypes of tolerance and sugar fermentation.

### Case Study 7

7.1 *Candida albicans* is a normal part of the human commensal flora, however it is also one of the most common fungal species that can cause human disease. Several cross-azole tolerance *C. albicans* strains were obtained by evolving with posaconazole from wild type diploid strain. (11)

Based on our method, trisomy of several chromosomes and LOH events is confirmed as previous report. Only dozens of high confidence somatic variations were detected. (Figure S2) Some genes with mutations more possibly related to azole tolerance have been manual picked up, including *ERG24*, *HMG1*, CAALFM_CR01280CA (sterol metabolic or transporter genes) and *PSY2*, *ALR1*, *MSS11*, CAALFM_C102280CA, CAALFM_C503940CA (genes with different mutations in independently evolved strains). (Table S3) The potential functions of these genes for azole tolerance are worth further research.

7.2 In another report about global analysis of mutations driving microevolution of diploid *C. albicans*, whole-genome sequencing was performed on multiple *C. albicans* isolates passaged both in vitro and in vivo to characterize the complete spectrum of mutations arising in laboratory culture and in the mammalian host. Microevolutionary patterns including de novo base substitutions and frequence of loss-of-heterozygosity events were reported.(12)

According to our analysis, most samples showed hundreds of high confidence somatic variations as new mutations. But one sample (P76055_GI_A, SRR5133898) showed hundreds of thousands of new mutations, which is unexpected, as so many mutations should not occur in the short-term microevolution processes theoretically. It is speculated that this sample might be the result of contamination during evolution experiments or genome sequencing, if this is not a mistake during uploading date to database. The microevolutionary patterns reported in this article might need to be re-evaluated. (Figure S2)

### Case Study 8

An extremely halotolerant fungi *Hortaea werneckii* with highly heterozygous genome in length of 50Mb was evolute for over seven years through at least 800 generations in a medium containing 4.3 mol/L NaCl. (13) Genome sequencing was performed for the original and the evolved strains, but variant calling was unsuccessful in the previous report (13). Based on our analysis, 22∼54 indels + 52∼126 SNVs were detected as high confidence somatic variations in the evolved strains. We manually pick out the mutation Gly160Asp in gene annotated as “basic leucine zipper (bZIP) domain of bZIP transcription factors”, which may be related to phenotype changes through global transcriptional regulation mechanisms. This mutation also appeared in another evolved strain EXF-225_evol110 (in KAI6820633.1, hypothetical protein KC358_g9374), indicating that this mutation might be related to halotolerance.

### Case Study 9

Finally, we also attempted to apply this approach to studies of prokaryotes. ATCC98092 is an *Escherichia coli* strain with a 4.5M chromosome and a 454kb plasmid. This is somewhat like multiple chromosomes.

The plasmid of ATCC98082 was cured in our study by CRISPR-Cas cutting. Then the analyce on the strains before and after the operation were performed using this method. CNV analysis results show that the plasmid has been cured, as expected. Only 2 high confidence somatic variations were detected, which should be the result of spontaneous mutations.

## Discussions

### Methodological Overview

We present a microbial comparative genomic analysis framework inspired by tumor-normal sequencing paradigms, which demonstrates robust performance in diploid and polyploid strain analysis while remaining applicable to haploid systems. This methodology is designed for researchers conducting high-resolution comparative genomic studies in microorganisms, particularly for investigations of laboratory evolution/microevolution, mutagenic breeding, spontaneous mutations, and related phenomena. Accurate mutation detection forms the cornerstone of downstream research and represents a critical focus. By systematically capturing evolutionary mutations (Table S1), this approach enables detailed retrospective analysis and facilitates advanced methodologies such as Genome-Wide Association Studies (GWAS). Integration with efficient microbial gene-editing technologies further allows mutation reconstruction and validation, thereby accelerating progress through the Design-Build-Test-Learn (DBTL) cycle in synthetic biology applications. (14)

Visualize chromosomal distributions of somatic variants, germline variants, and LOH patterns. Spatial clustering analysis differentiates localized LOH events from whole-chromosome alterations. VAF patterns provide ploidy estimation (e.g., 50% heterozygosity in diploids vs. 33% or 67% in triploids), complementing CNV data.

### Analytical Workflow

The framework comprises four principal components:

1. **Sample Designation:** Establish reference genome specifications and designate “normal” versus “tumor” samples for comparative analysis.
2. **Paired Sequencing Analysis:** Execute tumor-normal comparative sequencing to identify somatic mutations and loss of heterozygosity (LOH) events.
3. **Variation Visualization:** Generate chromosomal position-variant allele frequency (VAF) scatterplots to characterize mutation distributions.
4. **Copy Number Profiling:** Detect copy number variations (CNVs), including potential aneuploidy.

### Implementation Guidelines

- **Sample Selection:** Origin strain should serve as the normal sample, sequenced alongside experimental derivatives. Publicly available raw reads from equivalent strains may substitute to reduce costs, though disparities between “equivalent” strains require careful evaluation.(15)
- **Reference Genome Sequence:** Origin strain should serve as reference genome, ideally telomere-to-telomere assemblies from third-generation sequencing (16, 17). Publicly available reference genome from equivalent strains may also substitute to reduce costs and time. While draft assemblies are acceptable, completed assemblies are preferred. The genomes sequences of a strain from the same species are feasible but discouraged due to phylogenetic noise.

### Case-Based Validation

Several typical cases for researchers to troubleshoot (Figure 2):

a. Evolved trains with somatic mutations and LOH;
b. Evolved strains with somatic mutations, LOH, and aneuploid CNVs;
c. Haploid/homozygous derivatives showing complete LOH conversion;
d. Strains with distant genetic relationships manifesting as implausible variations (e.g. unexpected contamination during evolution experiments).

### Perspectives and Future Directions

While our tumor-normal paradigm represents a core approach, alternative microbial-adapted pipelines remain viable. Critical evaluation should prioritize compatibility with microbial genome architectures, which range from contig-level drafts to complete assemblies. Recent antifungal resistance studies in *Candida* species have successfully implemented analogous normal-referenced mutation detection via Mutect2. (18)

Current adaptations explore single-cell microbial sequencing to resolve genotype dynamics during evolution. The framework shows translational potential for clonal organisms in plant biology and lower animal research.

## Materials and Methods

### Comparative genomic analysis

Bwa 0.7.17-r1188 was used to map the reads to the reference genome sequences(19, 20). Then, “VarScan somatic” pipeline using samtools, picard and VarScan2 was performed with default parameters using “normal” control sample and “tumor” samples(21–23). The “germline variants” should be the ative heterozygous sites in the diploid reference genome and could also be used for removing false positives from the sample analysis. The “somatic variants” should be the mutations introduced by a genetic engineering process or generated during the evolution. The loss of heterozygosity (LOH) sites could also provide some hints for the genetic events during evolution. The duplications and deletions as well as possible aneuploidy in the evolved genome were identified by analyzing the coverage maps on the yeast chromosome, which were generated by “VarScan cnv” and cnvkit(23, 24). R-script using ggplot2 package were used for creating a scatter plot to present the distribution of variations.

The raw reads sequences and reference genome sequences are available as listed in Table S2

### Reference genome sequences

To obtain the high-quality reference genomes sequence of *Saccharomyces cerevisiae* Ethanol Red, ONT long reads (SRR15831142) were assembled by the Canu program(25). The assembled contigs were were order to chromosome sequences, and polished by pilon1.22 with short reads (SRR15831144) (26). Finally, the 12-Mb high quality genome sequences with 16 linear chromosomes, one circular mitochondrial and 7 unplaced contigs was obtained (GWHFGOL00000000)

The scaffold sequences of Candida albicans P76055 (AJJD01000000) were order to chromosome sequences (GWHFGOI00000000), by comparing to Candida albicans SC5314 chromosome sequences (GCF_000182965.3) as reference.

The scaffold sequences of Hortaea werneckii EXF-2000(GCA_000410955.1) were polished by GenomicConsensus(Quiver)2.3.3 with PacBio long reads (SRR5086620, SRR5086623, SRR5086626, SRR5086627 and SRR5086628) and NextPolish with short reads (SRR866616) (27). Finally, the 50-Mb corrected genome sequence was obtained (GWHFGOH00000000).

Annotations of reference genome sequences were performed by Funannotate 1.8.16, if necessary.(28)

## Authors’ contributions

SY conceived the project. JJY analyzed the data and performed the bioinformatic analyses. SY and JJY discussed the data and wrote the manuscript. All authors read and approved the final manuscript.

## Acknowledgments

This work was supported by National Natural Science Foundation of China (grant nos. 32071420, 21825804 and 31921006)

## Supplementary Materials

**Table S1. Sequencing data and reference genome details for case studies.**

**Table S2. Comprehensive analysis results of case studies.**

**Table S3. Putative azole tolerance-associated mutations identified in Case 6.**

**Figure S1.** Genome-wide distribution profiles of genetic variations (left) and copy number variations (CNVs, right).

(a) *ISO12*: Diploid strain exhibiting somatic variations, germline variations, loss of heterozygosity (LOH) on chromosome 10, and amplifications on chromosomes 4 and 10.

(b–d) *ScY01, ScY033T, and ScY001T*: Strains displaying somatic variations, germline variations, and LOH on chromosome 10, along with CNV on chromosome 9 leading to aneuploidy 2n+1(IX).

**Figure S2.** Genome-wide distribution profiles of genetic variations (left) and copy number variations (CNVs, right).

(a) *Kluyveromyces marxianus* KM-100d: Displays a ∼100 kb amplification region on chromosome 3 (CP015056.1: 1,476,415–1,571,769).

(b) *Candida tropicalis* X-17.2b and (c) *Candida albicans* EVO_P76055-2: Demonstrate characteristic diploid genomic profiles.

(d) *C. albicans* P76055_GI_A: Reveals >100,000 high-confidence somatic variations when compared to the normal sample *C. albicanhhs* P76055.

**Figure S3.** Venn diagram of high-confidence somatic variations shared across strains in Case 1(a), Case2(b) and Case 3(c).

