## Supplementary material for "Advancing Microbial Comparative Genomics Through Tumor-Normal Inspired Framework": Table S2

**Table S2 Comprehensive analysis results of case studies**

| **Case** | **Description** | **Confirmed** | **Revised** | **New discovery** | **Note** |
| --- | --- | --- | --- | --- | --- |
| **1** | a diploid *Saccharomyces cerevisiae* strain that overexpresses xylose catabolism genes under conditions of mixed xylose/glucose medium supplemented with sodium salts. The fully evolved yeast strains demonstrated efficient conversion of glucose and xylose in lignocellulose hydrolysates to ethanol, highlighting their potential for industrial applications. To uncover the genomic basis of these enhanced capabilities, we sequenced the final evolved strains, several intermediate strains during adaptive evolution and the original strain. |  |  | 288 high quality variations (231SNV + 57Indel)  Large LOH detected in chr4 and chr10 | Among these mutations, we discovered and experimentally confirmed that mutations in genes such as TRK1 and NFS1 are key phenotype related mutations. |
| **2** | an industrial *S. cerevisiae* that is strongly robust and has higher productivity, obtained by Genomewide Evolution-based CRISPR/Cas with Donor-free (GEbCD) system | Additional copy of chr9. | less variations than the previous report. | LOH event in several chromosomes | The effectiveness of this GEbCD system might need to be reevaluated and perhaps be more accurate than the previous report. |
| **3.1** | an evolved industrial strain, which can grow and ferment hexose sugars under a combination of stress factors (high temperature and lignocellulose-derived inhibitors) |  | less variations than the previous report. |  | Mutations, LOH and CNV may play a combined role in causing phenotypic changes |
| **3.2** | ScY01 derived from the industrial strain Ethanol Red through adaptive evolution at high temperature  ScY033T, thermotolerant strain  ScY001T, osmotolerant strain |  | less variations than the previous report. | Additional copy of chr9 in the strains. | Mutations, LOH and CNV may play a combined role in causing phenotypic changes in these strains. |
| **3.3** | Y01-tp, an haploid segregant from ScY01 |  |  | almost all of the variations are homozygous variations (LOH and homozygous somatic variation), | consistent with the characteristics of haploid.  This result provides a typical example for haploid (or highly homozygous) samples. |
| **4** | LSF1, another active dry yeast strain, reported from Lesaffre, but lacks detailed information | ploidy 3N+1(I) |  | Dozens of thousands of somatic variations were detected, with Ethanol Red as normal sample, | LSF1 should not be from Ethanol Red or close to Ethanol Red. This strain should be applied in fields such as Mantou or bread. |
| **5** | [Kluyveromyces marxianus developing ethanol tolerance during adaptive evolution with significant improvements of multiple pathways](https://www.ncbi.nlm.nih.gov/pubmed/30949239" \t "https://ngdc.cncb.ac.cn/gsa/browse/_blank) | SNV located in the coding region of SAN1, YAP1 etc |  | ~100kb amplification region, located at CP015056.1 (chr3) (1476415~1571769), | The amplified regain harboring several transcription related genes, such as IMP2 (sugar utilization regulatory protein IMP2), RRD1 (serine/threonine-protein phosphatase 2A activator 1), SLN1 (osmosensing histidine protein kinase SLN1), whose dose changes may be associated with ethanol tolerance. |
| **6** | Candida tropicalis adapted strain with increased tolerance to ethanol, furfural and hydroxymethylfurfural at high temperatures and improvement in fermentation ability at high glucose concentrations and xylose-fermenting ability。 |  |  | 373 high confidence somatic variations (163 SNV+210 Indel) | the mutations in the upstream region of NGT1(High-affinity glucose transporter 1) and GSF2 (Glucose signaling factor 2) related to the transcriptional changes of NGT1 and GSF2, and more genes regulated by them, affecting the phenotypes of tolerance and sugar fermentation. |
| **7.1** | Several cross-azole tolerance Candida  albicans strains were obtained by evolving with posaconazole | Trisomy of at least one of chromosomes |  | Only dozens of high confidence somatic variations were detected in these strains. | Some genes with mutations more possibly related to azole tolerance have been manual picked up, including ERG24, HMG1, CAALFM_CR01280CA (sterol metabolic or transporter genes) and PSY2, ALR1, MSS11, CAALFM_C102280CA, CAALFM_C503940CA (genes with different mutations in independently evolved strains). |
| **7.2** | Global analysis of mutations driving microevolution of diploid Candida albicans | CNV and LOH during evolution |  | Sample (P76055_GI_A, SRR5133898) showed hundreds of thousands new mutations, which is unexpected, as so many mutations should not occur in the short term theoretically. | It is speculated that this sample might be the result of contamination during evolution experiments or genome sequencing, if this is not a mistake during uploading date to database. The microevolutionary patterns reported in this article might need to be re-evaluated. |
| **8** | An extremely halotolerant fungi black yeast Hortaea werneckii was evolute for over seven years through at least 800 generations in a medium containing 4.3 M NaCl |  |  | ~50 indel + ~100 SNV was detected as high confidence somatic variations in the evolved strains. | Gly160Asp in gene annotated as basic leucine zipper (bZIP) domain of bZIP transcription factors, may be related to phenotype changes through global transcriptional regulation mechanisms. |
| **9** | Ecoli ATCC98082 cured the large plasmid |  |  | CNV analysis shows that the plasmid has been cured, as expected. Only 2 high confidence somatic variations were detected | The 2 high confidence somatic variations should be the result of spontaneous mutations. |
