## Supplementary material for "Advancing Microbial Comparative Genomics Through Tumor-Normal Inspired Framework": Table S3

**Table S3 Putative azole tolerance-associated mutations identified in Case 6**

| Gene ID | Gene | Gene annotation | Muations | Note |
| --- | --- | --- | --- | --- |
| CAALFM_C209400CA | ERG24 | delta(14)-sterol reductase | upstream_gene_variant in SRR18942590 | sterol metabolic |
| CAALFM_C103780CA | HMG1 | hydroxymethylglutaryl-CoA reductase (NADPH) | upstream_gene_variant in SRR18942591 | sterol metabolic |
| CAALFM_CR01280CA |  | sterol transporter | upstream_gene_variant in SRR18942596 | sterol transporter |
| CAALFM_C102280CA | PSY2 | Putative protein phosphatase PP4 complex subunit; macrophage-induced gene | upstream_gene_variant in SRR18942588 and SRR18942591 | different mutations in independently evolved strains |
| CAALFM_C302370CA | ALR1 | Mg(2+) transporter | upstream_gene_variant in SRR18942588 and SRR18942589 | different mutations in independently evolved strains |
| CAALFM_C503940CA |  | Putative multidrug resistance protein; upregulated by Efg1p | upstream_gene_variant in SRR18942593 and SRR18942596 | different mutations in independently evolved strains |
| CAALFM_CR04840CA | MSS11 | Transcription factor Mss11p | missense_variant p.Arg268His SRR18942591 and upstream_gene_variant in SRR18942596 | different mutations in independently evolved strains |
| CAALFM_CR02120CA | ECM4 | C2H2 zinc finger transcription factor | upstream_gene_variant in SRR18942589, SRR18942593 and SRR18942595 | different mutations in independently evolved strains |
