## Supplementary figures and images for "Advancing Microbial Comparative Genomics Through Tumor-Normal Inspired Framework"

### Figure S1

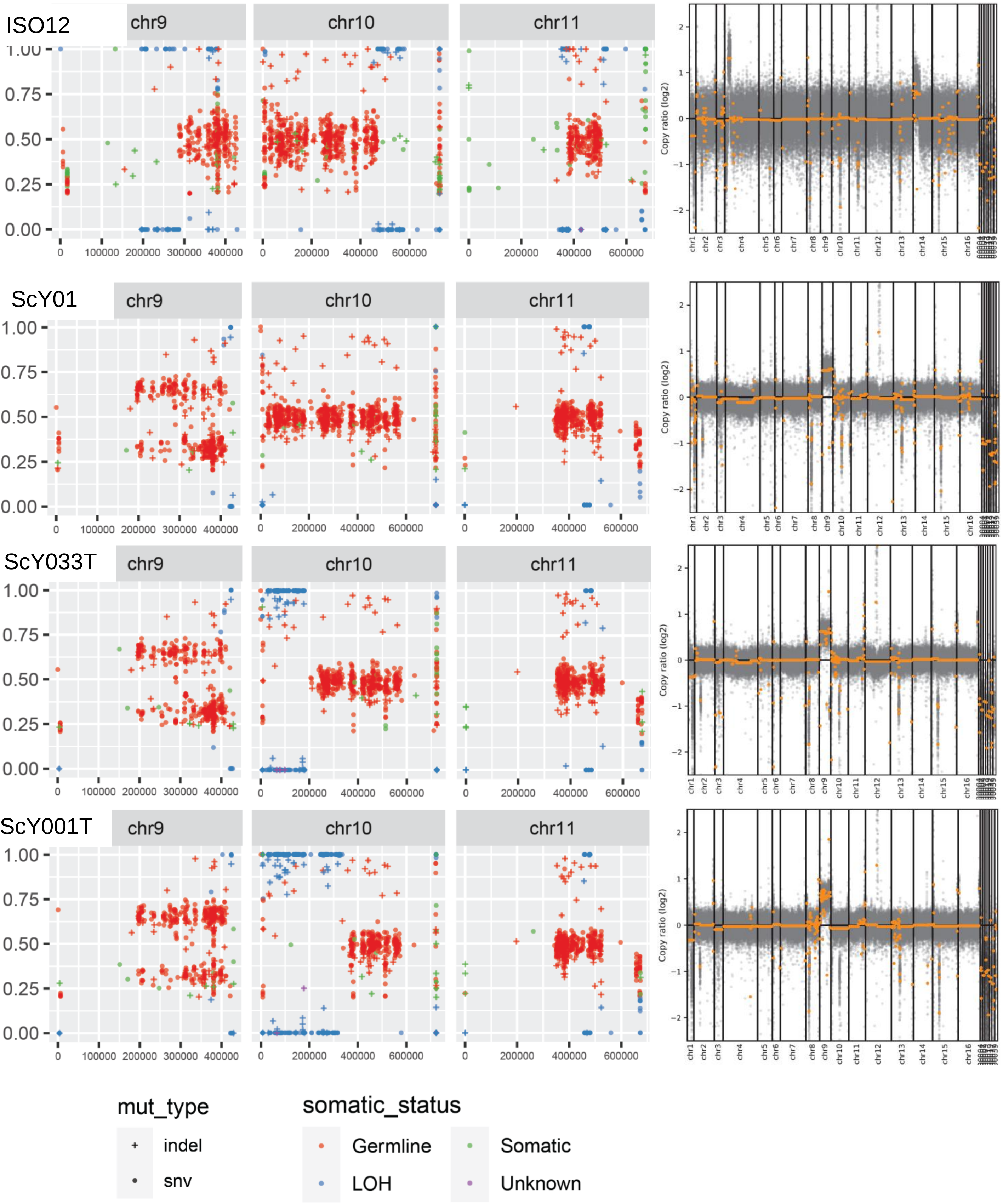

### Figure S2

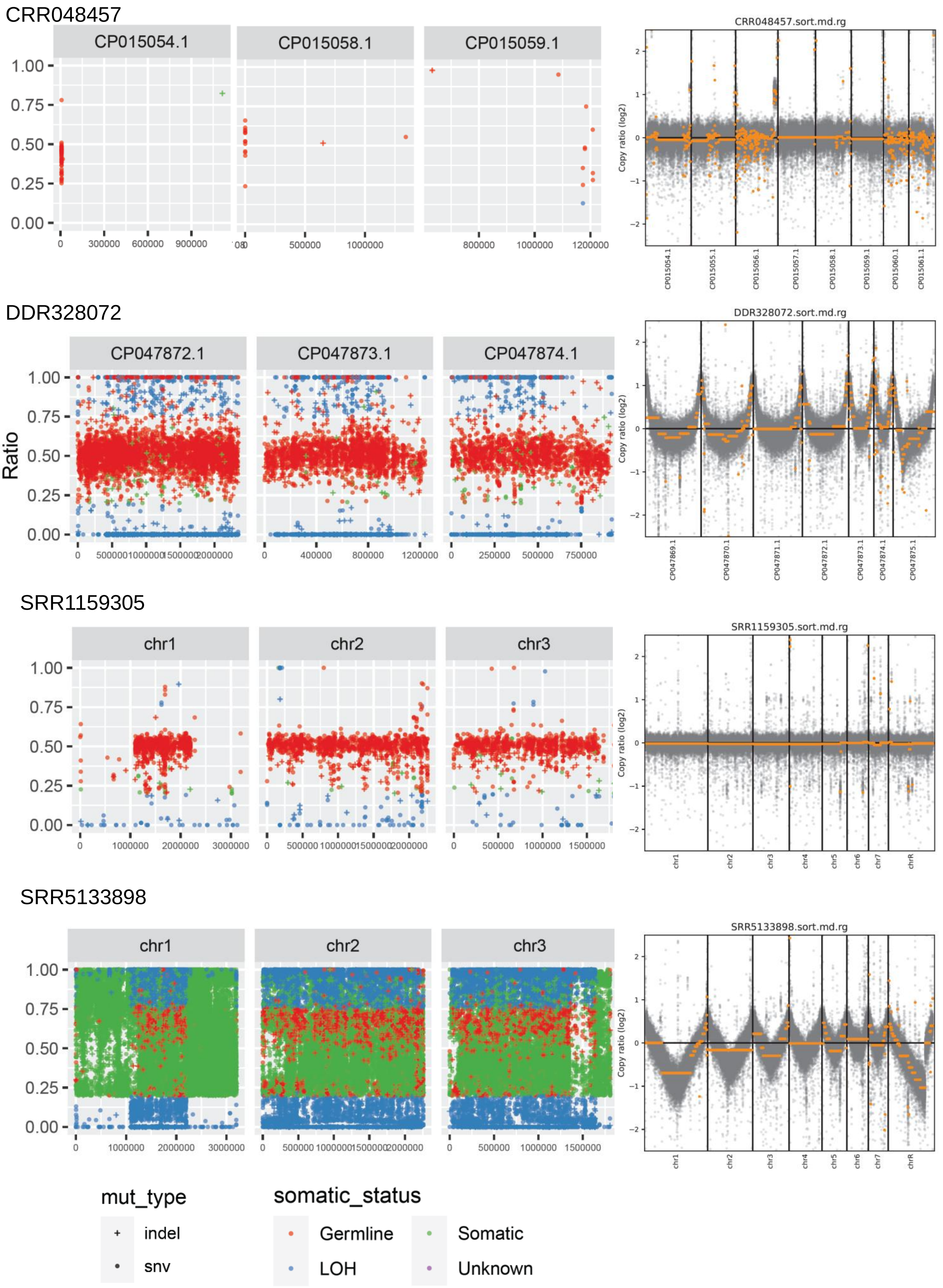

### Figure S3

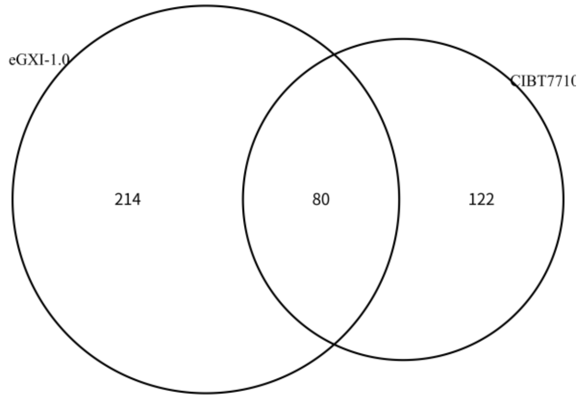

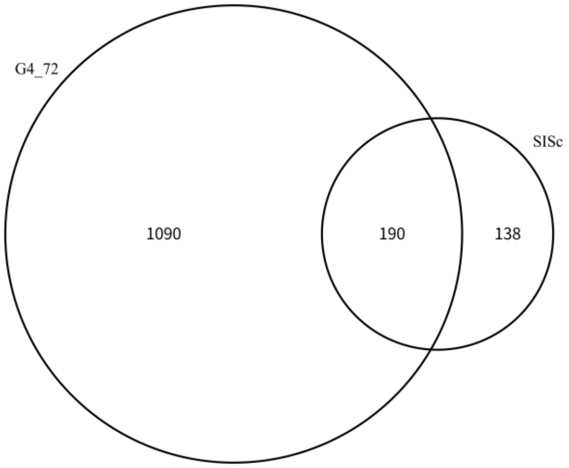

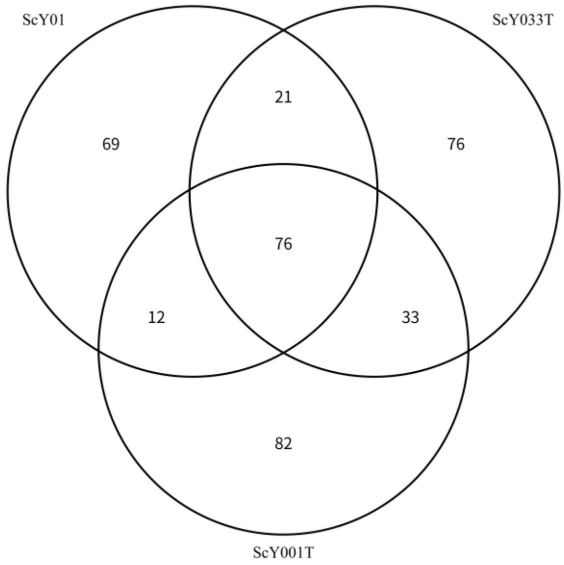
